# Apigenin relaxes rat intrarenal arteries: involvement of Cl^−^ channels and K^+^ channels

**DOI:** 10.1101/371849

**Authors:** Yixin Jing, Miaomiao Dong, Yu Liu, Xiaomin Hou, Pengmei Guo, Weiping Li, Mingsheng Zhang, Jiyuan Lv

**Affiliations:** The First Clinical Hospital, Shanxi Medical University, Taiyuan 030001, Shanxi Province, China; Department of Pharmacology, Shanxi Medical University, Xinjiannanlu 56, Taiyuan 030001, Shanxi Province, China

**Keywords:** apigenin, intrarenal artery, vasorelaxation, flavonoid, calcium-activated chloride channels, voltage-dependent potassium channels, inwardly rectifier potassium channels

## Abstract

The vasodilator effect of apigenin (API) was demonstrated in a number of vascular beds. We aimed to characterize the vasospasmolytic and electrophysiological effects of apigenin (API) in intrarenal arteries (IRAs). The vascular tone of male rat isolated IRAs was recorded with a myograph. Transmembrane Cl^−^ currents through Ca^2+^-activated Cl^−^ channels (CaCCs), K^+^ currents through voltage-gated K^+^ (Kv) channels and inwardly rectifier K^+^ (Kir) channels were recorded with patch clamp in the freshly isolated arterial smooth muscle cells (ASMCs). Preincubation with API (10-100 μM) concentration-dependently depressed the contractions induced by KCl, 9,11-dideoxy-9α,11α-methanoepoxy prostaglandin F_2α_ (U46619), phenylephrine and vasopressin without significant preference and the IC_50_ values were 13.27-26.26 μM. Acute application of API elicited instant relaxations in the IRAs precontracted with these vasoconstrictors and the RC_50_ values were 5.80-24.33 μM. API relaxation was attenuated by chloride deprivation, CaCC blockers, Kv blocker and nitric oxide synthase inhibitor, but not by Kir blocker and cyclooxygenase inhibitor. At 10-100 μM, API depressed CaCC currents and Kir currents while enhanced Kv currents of IRA ASMCs. The present results demonstrate that API counteracts various vasoconstrictors noncompetitively and nonspecifically and suggest that modulation of CaCCs, Kv and Kir channels of IRA ASMCs is involved in its vasospasmolytic effects.

## INTRODUCTION

Apigenin (API, C_15_H_10_O_5_, MW270.24), a natural flavonoid present in a lot of edible plants such as celery, parsley and oranges, possesses significant pharmacological activities and has been suggested to treat a variety of disorders such as cancers [1], neurodegeneration [2], metabolic syndrome [3], hyperlipidemia [4], vasodilative impairments [5-8] and vascular restenosis [9]. API-induced vasodilation was demonstrated in rat aorta [10-13], pial artery [14] and mesenteric artery [15]. The suggested mechanisms underlying its vasodilation include production of nitric oxide and guanosine 3’, 5’-cyclic monophosphate, activation of transient receptor potential vanilloid 4 cation channel, calcium-activated potassium channels (K_Ca_) and ATP-sensitive potassium channels (K_ATP_), inhibition of extracellular Ca^2+^ influx. Review and analysis of available reported researches point to that API may have multiple targets within vascular smooth muscle cells (VSMCs) and may be classified as a sort of pleiotropic drug. In this case, the exact mechanisms underlying its vasoactive actions need further deep investigations.

The K^+^ channels are very important regulators of the myocyte membrane potential, vascular resistance and eventually blood flow [16,17]. Most commonly expressed K^+^ channels in VSMCs are voltage-dependent K^+^ channels (Kv), inwardly rectifier K^+^ channels (Kir), K_Ca_ and K_ATP_. Kv channels were proved to be involved in renal arterial tone regulation [18-20]. The existence and importance of Kir channels were also demonstrated in intrarenal arterioles (IRAs) [21,22]. Recent studies showed that calcium-activated Cl^-^ channels (CaCCs) play a very important role in vascular tone modulation [23]. To our knowledge, the vasomotor effects of API on IRAs and electrophysiological effects of API on Cl^-^ channels, Kv channels and Kir channels of arterial smooth muscle cells (ASMCs) of IRAs have not been addressed. The present experiments were designed to obtain a clearer insight into the effects of API on IRAs and a deeper understanding on the underlying mechanisms.

## MATERIALS and METHODS

### Animals

Adult healthy male Sprague-Dawley rats (body weight: 250-300g) provided by Animal Center of Shanxi Medical University, China. All protocols and procedures of this study were approved by the Animal Care and Use Committee of Shanxi Medical University and conform to NIH guidelines for the care and use of laboratory animals. The second and third orders of intrarenal arteries (IRAs, inner diameter: 220-320 μm) were gently isolated for myograph and patch clamp study, after anesthesia with intraperitoneal administration of sodium pentobarbital (40 mg/kg) and euthanasia.

### Drugs and Chemicals

Apigenin (API, HPLC>98%) was purchased from PERFEMIKER (Shanghai Canspec Scientific Instruments Co). 4-(2-hydroxyethyl) piperazine-1-ethanesulfonic acid (HEPES), ethyleneglycol-bis(β-aminoethylether)-N,N,N’,N’-tetraacetic acid (EGTA), 9, 11- dideoxy-9α, 11α-methanoepoxy prostaglandin F2α (U46619), phenylephrine (PE), vasopressin (VP),4-aminopyridine (4-AP), NG-nitro-L-argininemethylester ester (L-NAME), tetreathylammonium(TEA), niflumic acid (NFA), CaCC_inh_-A01, papain, indomethacin, bovine serum albumin, collagenase F, collagenase H and dithiothreitol were purchased from Sigma (St. Louis, MO, USA). API and indomethacin were dissolved in dimethylsulfoxide (DMSO) respectively just before use. When DMSO was used as solvent, its final concentration in the bath was less than 0.1%. All other reagents were dissolved in distilled water just before use.

### Measurements of arterial tension and tissue bath solutions

Isolation, mounting, vessel tone normalization, tissue bath solutions and general protocols were same as previously described [24] except illustrated elsewhere. Briefly, the cylindrical IRA rings (2 mm-long) were threaded with two 25-μm-diameter stainless steel wires and mounted transversely on a wire myograph. The vessels were bathed in a chamber containing 5 ml of physiological saline solution (PSS) and stretched gently to opposite directions to produce a tone roughly equivalent to 80 mmHg. The normal PSS composed of (mM) 118 NaCl, 4.7 KCl, 1.2 KH_2_PO_4_, 1.2 MgCl_2_•H_2_O, 20 NaHCO_3_, 10 HEPES, 2.5 CaCl_2_ and was bubbled with 95% O_2_ + 5% CO_2_ at 37 °C, pH=7.4. In Cl^-^-free bath solution, NaCl, KCl, CaCl_2_ and MgCl_2_ were replaced with equal moles of sodium D-gluconate, potassium D-gluconate, calcium gluconate and magnesium sulfate respectively.

### Cell isolation and patch clamp study

General procedures of isolation of single IRA ASMCs, preparation of electrodes, current recording and data analysis were same as previously described [24] except illustrated elsewhere. All currents were normalized with cell capacitance, expressed in pA/pF and recorded before (control), during the presence of API and after washout of API.

The chloride currents through CaCCs were recorded as reported methods [25], the bath solution contained (mM): 135 NaCl, 5.4 CsCl, 1 MgCl_2_, 1 CaCl_2_, 0.33 NaH_2_PO_4_, 5 TEA-Cl, 10 HEPES and 10 glucose (pH 7.35 adjusted with NaOH). The pipette solution contained (mM): 110 CsCl, 20 TEA-Cl, 2 MgATP, 10 EGTA, 5 HEPES and 0.16 MgCl_2_ (pH 7.2 adjusted with CsOH). CaCl_2_ 7.475 mM was also included to set the free Ca^2+^ concentration at 500 nM. Contamination by other currents was minimized by replacing K^+^ ions with Cs^+^ and by adding TEA chloride in both pipette and bath solutions. Cell were held at holding potentials of-100 mV and subjected to step depolarizations of 500 ms to +100 mV in 10 mV increments.

To record Kv current selectively, extracellular Ca^2+^ was deprived from the cell bath solution and high concentration of ATP and EGTA were included in the pipette solution to minimize K_ATP_ and K_Ca_ currents [26]. The pipette solution consisted of (mM) 110 KCl, 1.2 MgCl_2_, 5 Na_2_ATP, 10 HEPES, 10 EGTA, pH adjusted to 7.4 with KOH [26]. In this condition, the remainder currents recorded were markedly reduced by application of specific Kv blocker 4-AP (3 mM) [27]. Cells were held at holding potentials of −60 mV and subjected to step depolarizations of 500 ms to +60 mV in 10 mV increments. Kv currents were recorded and current-voltage (I-V) curves were plotted with the readings at the end of the pulse.

In recording Kir current, the cell bath solution contained (mM): 140 NaCl, 5.4 KCl, 0.33 NaH_2_PO_4_, 1.0 MgCl_2_, 1.8 CaCl_2_, 0.5 CdCl_2_, 5.0 HEPES, 10 glucose [28,29] and the pipette solution contained (mM): 100 potassium gluconate, 30 KCl, 5 EGTA, 5 HEPES, 1 MgCl_2_, 1 Mg-ATP, 3 K_2_-ATP [28,29]. Both bath and pipette solution contained 4-AP (3 mM), TEA (3 mM), glibenclamide (1 μM) and nifedipine (1 μM) in order to exclude transmembrane currents via Kv, K_Ca_, K_ATP_, Ca^2+^ currents and other Ca^2+^-activated currents [29-32]. Cells were held at −60 mV and subjected to stepwise test potentials from −160 mV to 0 mV in 10 mV increments for 500 ms at each potential [29,33]. Elicited currents were filtered at 1~2 kHz, averaged during the last 200 ms of each step, normalized to cell capacitance and used for construction of I-V curves.

### Data analysis

Results are means ± SEM of *n* IRA rings or cells. Each ring or cell was isolated from a separate animal. Paired or unpaired Student’s t-test was used to analyze data of two groups. Two-way analysis of variance (ANOVA) was performed for data of more than two groups. Differences were considered statistically significant when P<0.05. The values of RC_50_ (vasodilator concentration needed to decline the precontraction by 50%), IC_50_ (antagonist concentration needed to depress the maximal contraction by 50%) and EC_50_ (agonist concentration needed to produce 50% of the maximal contraction) were calculated by non-linear regression with GraphPad Prism ®, version 6.00 (GraphPad Software, San Diego, CA, USA).

## RESULTS

### Effects of preincubation with API on KCl-, U46619-, PE-, VP-induced contraction

KCl (20-108 mM, Fig.1A), U46619 (10^-8^-10^-5^ M, Fig.1B), PE (10^-7^-10^-5^ M, Fig.1C) and VP(10^-7^-10^-5^ M, Fig.1D) induced concentration-dependent contraction in IRA rings. The maximal contractions were 7.64 ± 3.08 mN, 10.63 ± 2.87 mN, 8.40 ± 2.10 mN and 9.19 ± 4.93 mN respectively; the values of EC_50_ were 33.49 ± 1.45 mM, 0.16 ± 1.57 μM, 0.51 ± 1.14 μM and 0.35 ± 1.25 μM respectively. Pretreatment with API shifted all of these concentration-contraction curves downwards and nonparallel to the right. At 100 μM, API depressed the maximal contractions by 81.15 ± 33.35%, 75.74 ± 24.22%, 84.05 ± 37.96% and 91.73 ± 23.98% for KCl,U46619, PE and VP, respectively. The IC_50_ values of API were 26.26 ± 1.09 μM, 18.19 ± 1.01μM, 22.77 ± 1.14 μM and 13.62 ± 1.44 μM, respectively.

**FIGURE 1.**
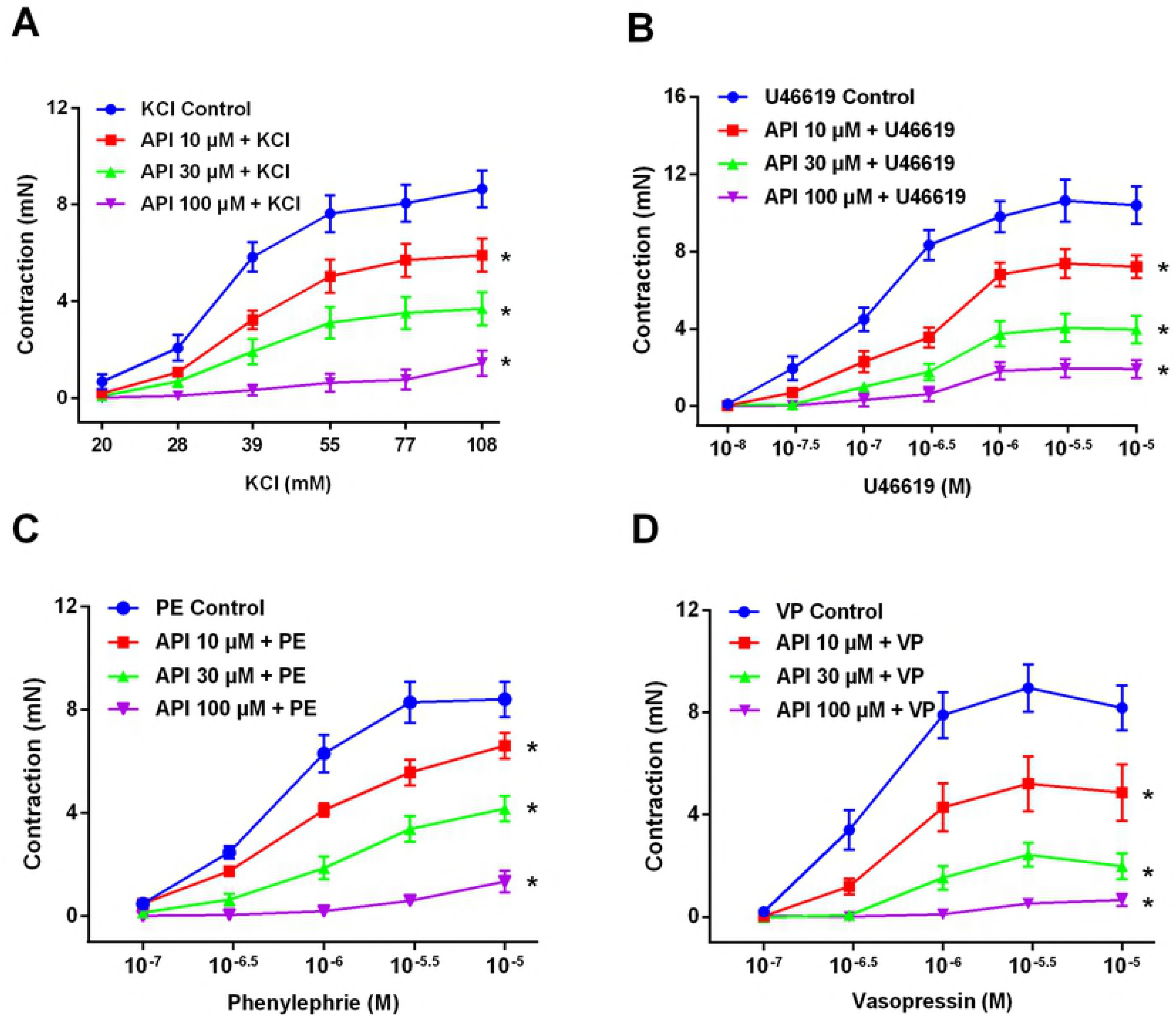
API depressed depolarization-, U46619-, PE-and VP-induced IRA contractions noncompetitively and nonspecifically. The contractive responses to the cumulative increment of KCl (A), U46619 (B), PE (C) or VP (D) were observed in the absence (control) or presence of API (10, 30, 100 μM) in isolated IRA rings. When the vessel tension recovered to the basal tone after the control concentration-contraction curve, API was added to the bath 20 minutes before the curves of KCl, U46619, PE or VP were reconstructed again. Contractions (mN) are presented as means ± SEM of 7 IRA rings isolated from 7 separate rats. * P<0.05 vs the respective control.

### Relaxation of API on the precontractions

To observe the direct vasorelaxation of API on IRAs, API (1-100 μM) was cumulatively added to the bath when the precontraction induced by KCl, U46619, PE or VP was sustained. Fig.2 showed that API concentration-dependently declined the precontractions (Fig. 2A) and the RC_50_ values were 24.33 ± 2.75 μM, 16.41 ± 1.16 μM, 17.89 ± 1.21 μM, 5.80 ± 1.45 μM for KCl-,U46619-, PE-and VP-induced precontraction (Fig. 2B and C), respectively. DMSO (vehicle) at up to 0.1% failed to affect the precontractions.

**FIGURE 2.**
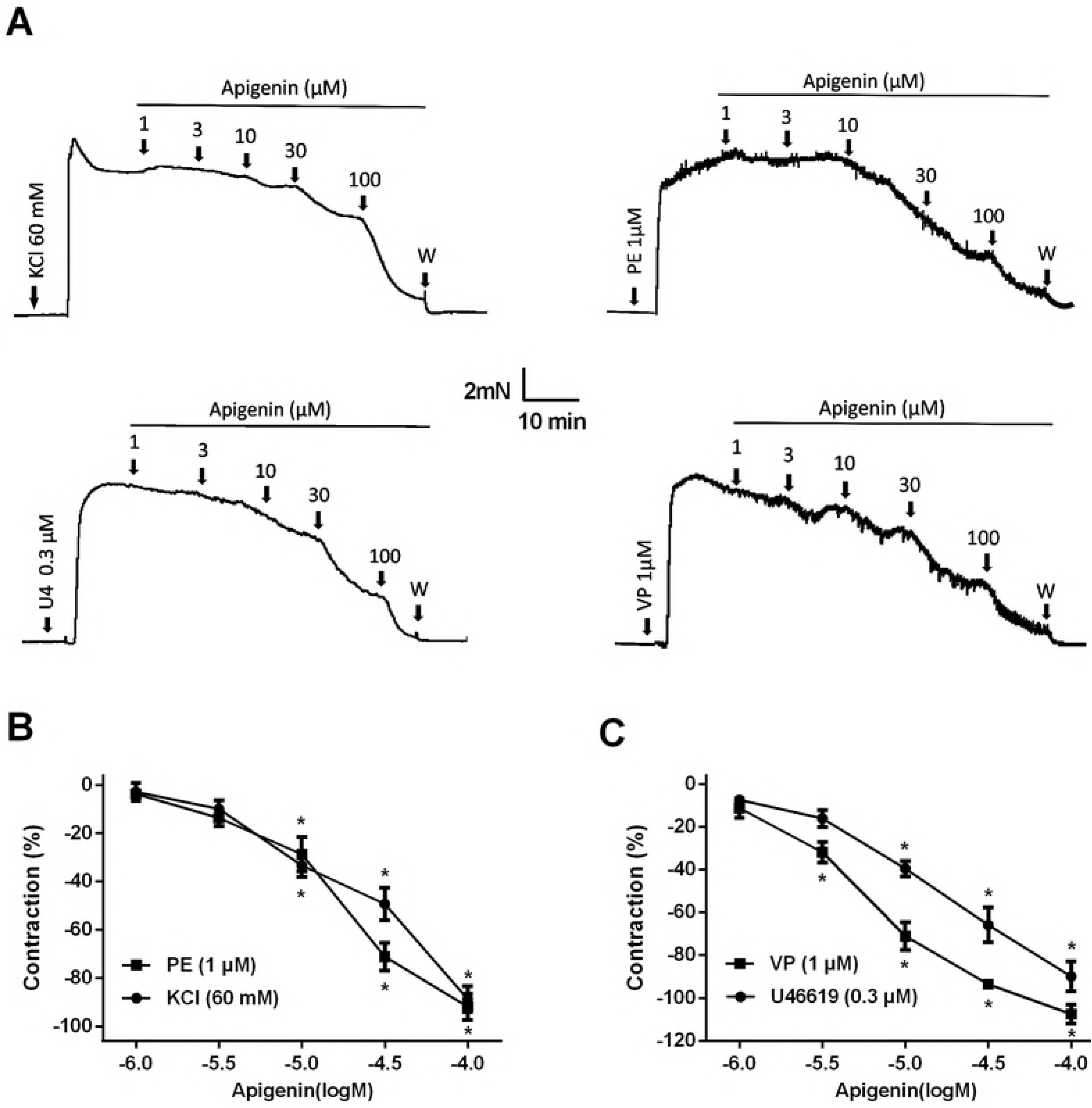
API relaxed IRA rings precontracted with KCl, U46619, PE or VP. Relaxations are expressed as percentages of the precontraction induced by KCl (60 mM), U46619 (0.3 μM), PE (1 μM) or VP (1 μM), respectively. A: Original recording of the vasorelaxations induced by cumulative addition of API on IRAs precontracted with KCl, U46619, PE or VP. B and C: Pooled data (means ± SEM) of API against KCl, PE (B), U46619 and VP (C), n=7. * P<0.05 vs vehicle (0.1% DMSO) control.

### Effects of various inhibitors on API-induced relaxation in IRAs

To explore the mechanisms underlying API-induced relaxation, inhibitor study was performed. As at 30 μM, API produced a solid and repeatable relaxation on the precontractions induced by various agonists, we chose this concentration to study effects of various inhibitors on the relaxation. Addition of certain inhibitors might produce a superimposed contraction upon KCl-induced or U46619-induced precontraction in some rings. If the superimposed contraction surpassed 10% of the original precontraction, the results were discarded. Preincubation with L-NAME (0.01mM), 4-AP (0.3 mM), NFA (3 μM) and CaCC_inh_-A01 (3 μM) reduced the API-induced relaxation by 34.40 ± 8.45%, 30.30 ± 9.26%, 39.62 ± 9.96% and 43.83 ± 9.39% upon 60 mM KCl-induced contraction (Fig.3A), and by 36.09 ± 9.08%, 33.84 ± 11.88%, 38.03 ± 8.94% and 43.49 ± 10.49% upon 0.3 μM U46619-induced contraction (Fig.3B). Neither Indo (0.01 mM) nor BaCl_2_ (30 μM) affected API-induced relaxation significantly.

**FIGURE 3.**
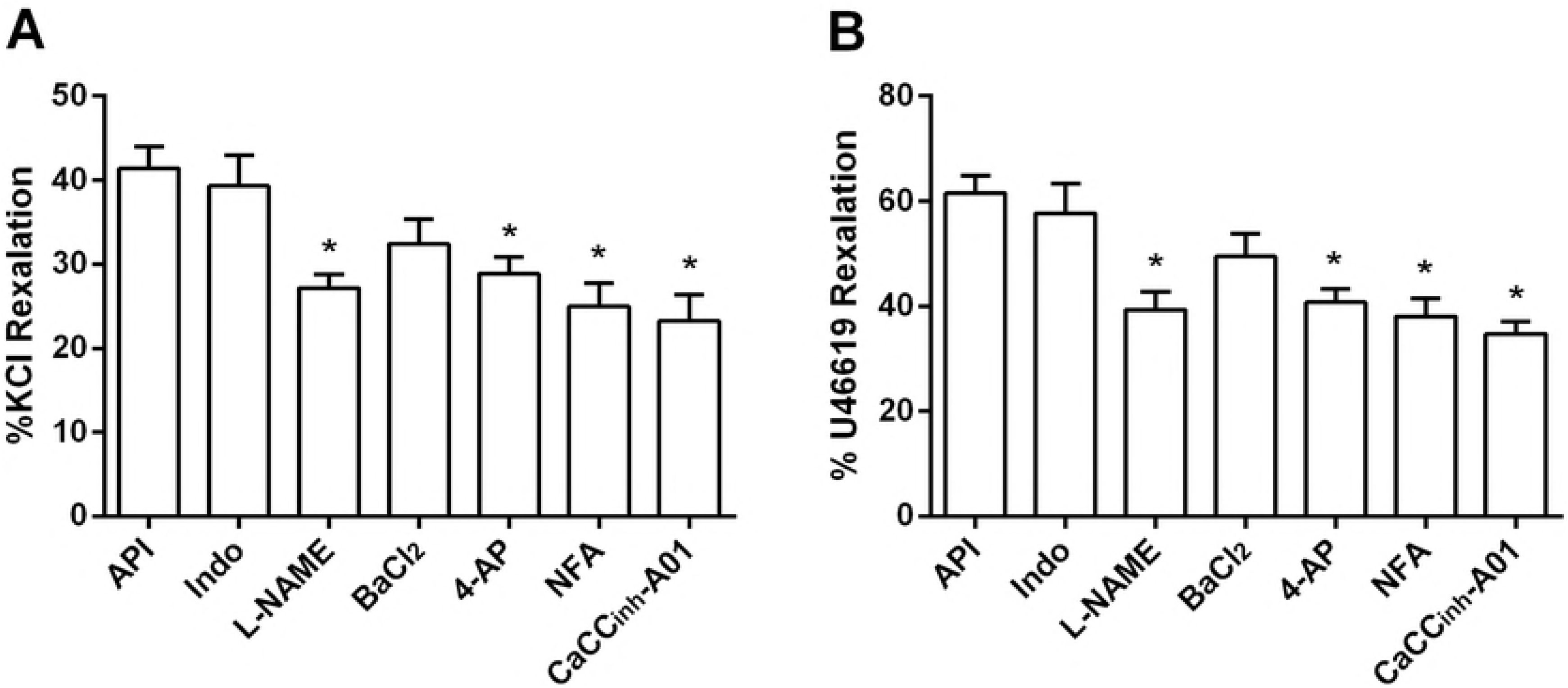
Effect of different inhibitors on API-induced relaxation in IRA rings precontracted by KCl or U46619. When the precontraction induced by either 60 mM KCl or 0.3 μM U46619 was sustained, Indo (0.01 mM), L-NAME (0.01 mM), 4-AP (0.3 mM), BaCl_2_ (30 μM), NFA (3 μM) or CaCC_inh_-A01 (3 μM) was added to the bath respectively. When the contraction was steady again in the presence of one of these inhibitors, API (30 μM) was added to bath. The API-induced vasodilations in the absence (control) or presence of inhibitors are expressed as percentage of the precontraction induced by KCl (A) or U46619 (B). n=7. * P<0.05 vs control.

### Effect of Cl^-^ deprivation on API-induced relaxation

The possible involvement of chloride channels in API-induced relaxation was also studied by observing the influence of Cl^-^ deprivation on the relaxation. Preincubation for 30 min with Cl^-^-deprived PSS solution, in which NaCl, KCl, CaCl_2_ and MgCl_2_ were replaced correspondingly with equal moles of sodium gluconate, potassium gluconate, calcium gluconate and magnesium gluconate, reduced 60 mM K^+^- and 0.3 μM U46619-induced contraction by 22.45 ± 10.56% and 28.14 ± 12.89% respectively. Fig.4 showed that compared with in normal PSS, API-induced relaxations on K^+^-contraction in Cl^-^-deprived PSS was reduced (52.15 ± 19.33% vs 32.81 ± 17.35%, P<0.05; API relaxation on U46619 contraction in Cl^-^-deprived PSS was reduced (58.92 ± 19.96% vs 39.31 ± 17.91%, P<0.05).

**FIGURE 4.**
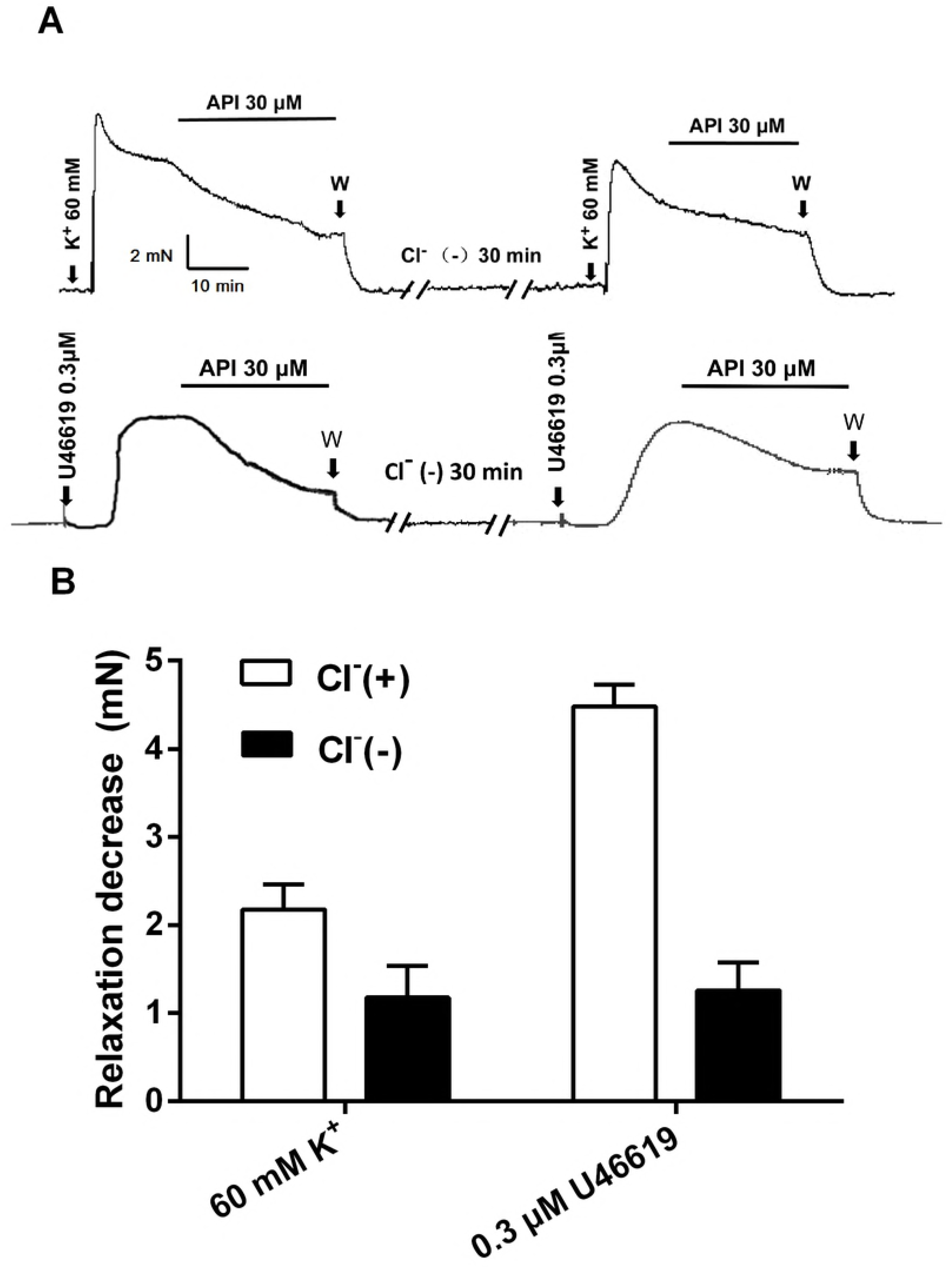
Cl^-^ deprivation attenuated API-induced relaxation in IRA precontracted with K^+^ or U46619. After API relaxations on the precontraction induced by KCl (60 mM) or U46619 (0.3 μM) were recorded in normal PSS solution (Cl^-^(+)), the vessels were washed with normal PSS solution and left alone to recover to the basal tone. Thereafter the vessels were incubated in Cl^-^ deprived PSS solution (Cl^-^(-)) for 30 min before next repetitive precontraction-relaxation procedure in Cl^-^-free solution. A: Original tracings of API-induced relaxation on K^+^- or U46619-induced precontraction in either normal or Cl^-^ deprived PSS solution. B: Pooled data of API-induced decline (percentage) on the precontractions. Results are means ± SEM. n=7. * P<0.05 vs Cl^-^(+).

### Effects of API on Cl^-^ currents of IRA ASMCs

The stable peak Cl^-^ current at a testing potential of +100 mV was 1338.52 ± 207.15 pA and current density was 53.54 ± 9.50 pA/pF (n=7). 4-AP (3 mM) suppressed voltage-dependent potassium channel current density by 72.01 ± 13.17%. API (10, 30, 100 μM) reduced the peak Cl^-^ current density by 17.96 ± 6.76%, 44.58 ± 12.57%, 65.11 ± 19.19% respectively (Fig. 5A and B). The action of API was fast (~20 s), relatively stable within 3 min and reversible upon washout of the drug (Fig. 5C).

**FIGURE 5.**
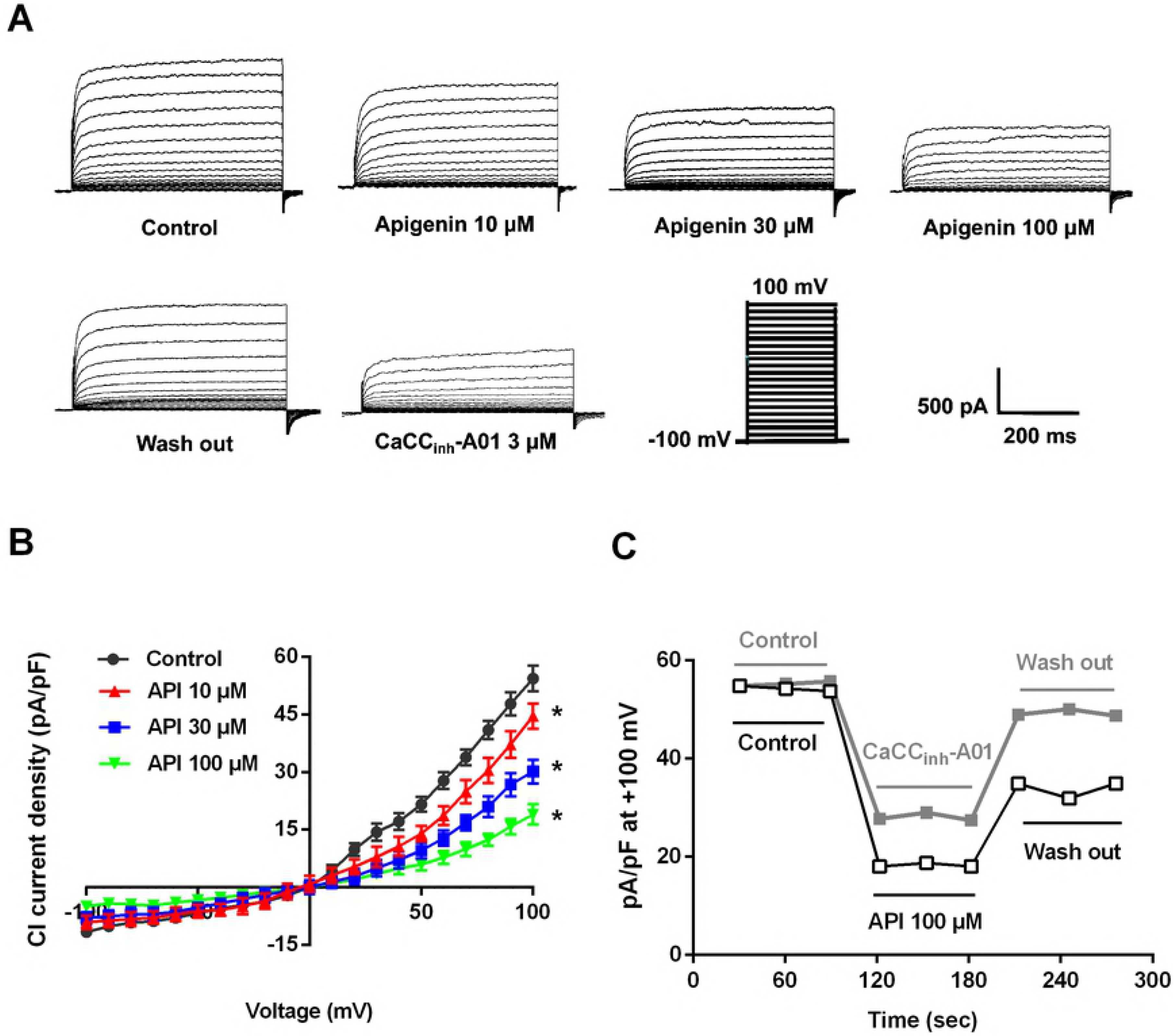
API inhibited calcium-activated chloride currents in freshly isolated single IRA ASMCs. A: Original recordings of the outward currents evoked by a series of depolarizing pulses (from −100 mV to +100 mV in 10-mV increments, 500 ms duration) in the absence (control) or presence of API (10, 30, 100 μM). B: Pooled I-V curves of CaCC currents. Data are presented as means ± SEM, n=7. *P<0.05 vs control. C: Diagrammatic time-course of CaCC currents recorded before (control), during the presence of API (100 μM) or CaCC_inh_-A01 (3 μM) and after washout of the drug at a testing potential of +100 mV.

### Effects of API on Kv currents in IRA ASMCs

The stable peak Kv current at a testing potential of +60 mV was 1256.89 ± 219.56 pA and current density was 43.34 ± 9.35 pA/pF (n=7). API (10, 30, 100 μM) increased the peak Kv current density by 28.56 ± 9.55%, 54.90 ± 9.49%, 81.61 ± 10.03% respectively (Fig.6 A and B). Again, the action of API on Kv currents was reversible (Fig. 6C).

**FIGURE 6.**
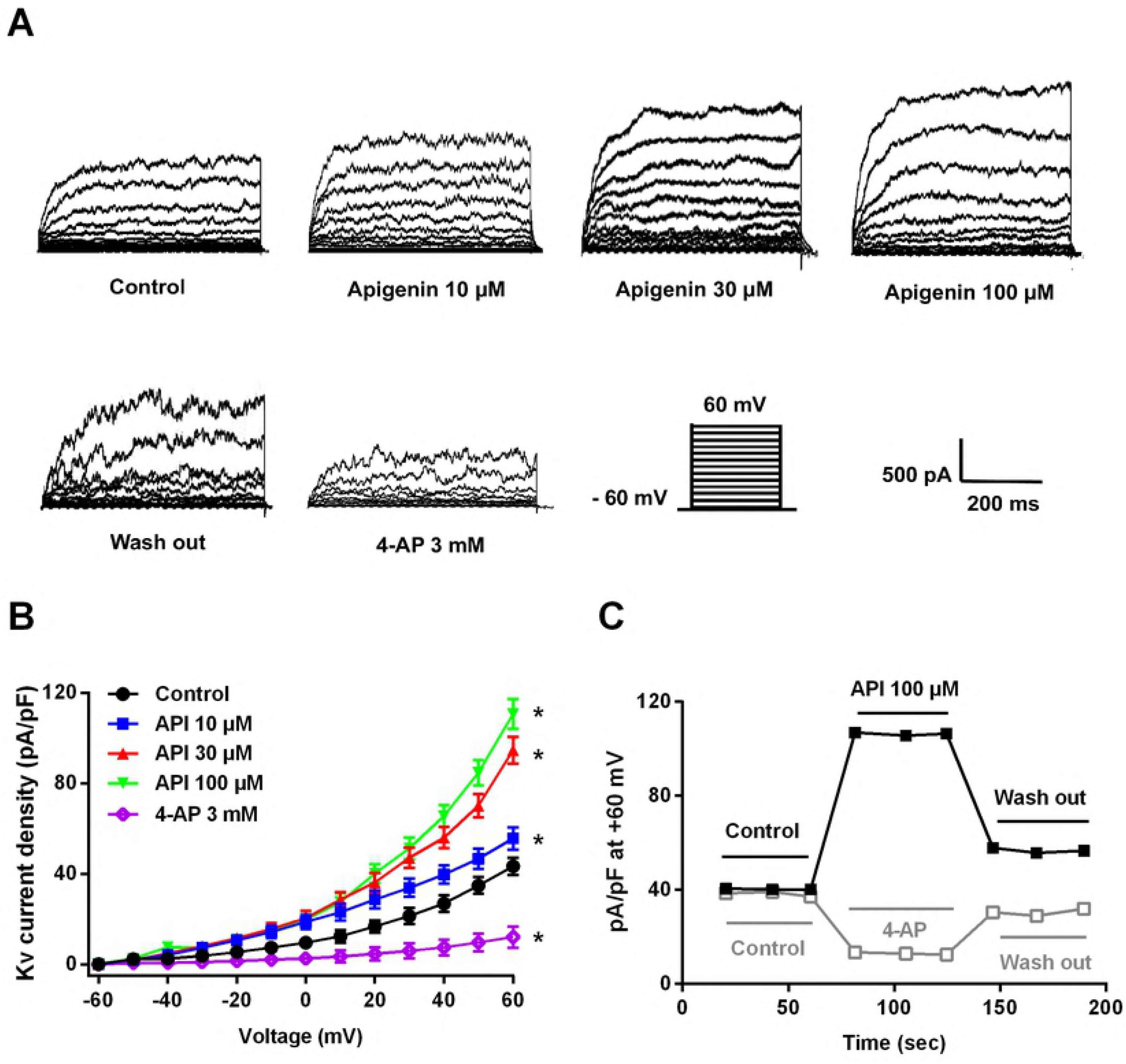
API enhanced Kv currents in freshly isolated single IRA ASMCs. A: Original recordings of the outward currents evoked by a series of depolarizing pulses (from −60 mV to +60 mV in 10 mv increments, 500 ms duration) in the absence of presence of API (10, 30, 100 μM). B: Pooled I-V curves of Kv currents in the absence (control) or presence of API. Data are presented as means ± SEM, n=7. *P<0.05 vs control. C: Diagrammatic time-course of Kv currents recorded before (control), during the presence of API (100 μM) or 4-AP (3 mM) and after washout of the drug at a testing potential of +60 mV.

### Effects of API on Kir currents in IRA ASMCs

The stable peak current of Kir at a testing potential of −160 mV was −313.24 ±77.43 pA and current density was −22.67 ± 5.16 pA/pF (n=7). API (10, 30, 100 μM) reduced the peak Kir current density by 12.57 ± 4.29%, 37.32 ± 8.37%, 46.63 ± 9.22%, respectively (Fig. 7A, C, D). Similar to its action on CaCC currents and Kv currents, the action of API on Kir currents was fast and reversible upon washout of the drug (Fig. 7B).

**FIGURE 7.**
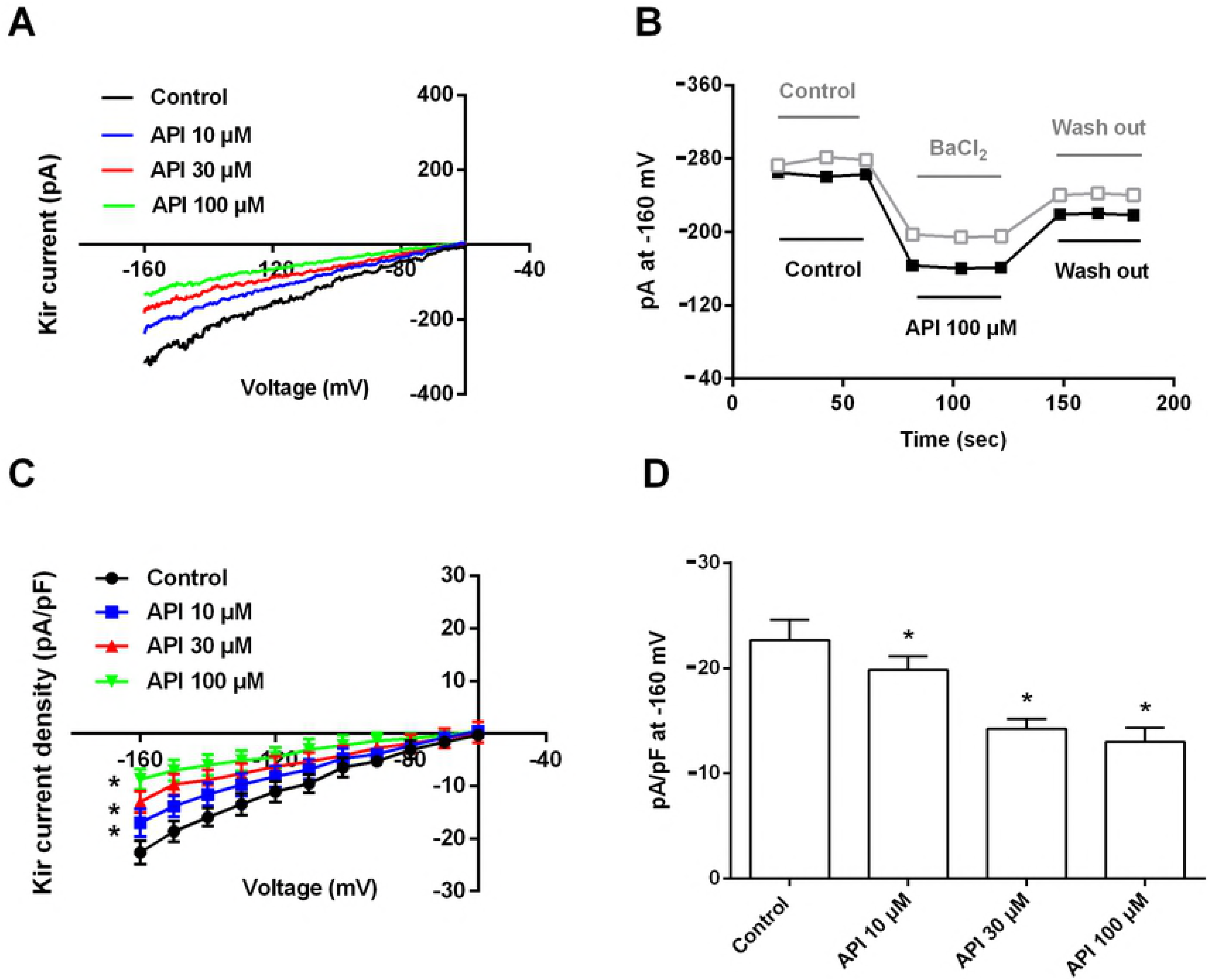
Effect of API on inward rectifier potassium channels in isolated rat IRA ASMCs. A: Original recordings of currents evoked by a series of depolarizing pulses (from −160 mV to 0 mV in 10-mV increments, 500 ms duration) in the absence (control) or presence of API (10, 30, 100 μM). B: Diagrammatic time-course of Kir currents recorded before (control), during the presence of API or BaCl_2_ (30 μM) and after washout of API at a testing potential of −160 mV. C: Summary of API effects on the I-V curves of Kir current density. D: API effect on Kir current density at −160 mV. Data are presented as means ± SEM, n=7. *P<0.05 vs control.

## DISCUSSION

The main findings of the present study are: 1. API was vasospasmolytic against various vasoconstrictors in IRAs. 2. API-induced IRA relaxation was attenuated by deprivation of extracellular chloride, CaCC blocker, Kv blocker and nitric oxide synthase inhibitor, but not by Kir blocker and cyclooxygenase inhibitor. 3. API depressed CaCC currents and Kir currents while enhanced Kv currents of IRA ASMCs.

The present study demonstrated that, preincubation with API, at concentrations reachable after oral administration [34,35], depressed depolarization-, U46619-, PE-and VP-induced IRA contractions and acute application of API of the same range of concentrations instantly declined the precontractions induced by these vasoconstrictors. Comparison of action potencies of API against these vasoconstrictors revealed the vasospasmolytic characteristics of API on IRAs, that is, API counteracts these vasoconstrictors concentration-dependently, noncompetitively and nonspecifically.

To explore the underlying mechanisms, inhibitor study was performed. API-induced vasorelaxation was significantly attenuated by nitric oxide synthase inhibitor L-NAME but not by cyclooxygenase inhibitor indomethacin, suggesting that nitric oxide production but not prostanoid production is involved in the vasorelaxation. This is consistent with the reported results that API vasorelaxation in rat pial arteries was attenuated by inhibition of nitric oxide synthase [14] and that API enhanced rat aortic endothelial nitric oxide synthase activity and endothelial nitric oxide synthesis [6]. API relaxation was also reduced by Cl^-^ deprivation, NFA, CaCC_inh_-A01and 4-AP, suggesting that chloride channels, Kv channels may be involved in the relaxation.

A variety of ion channels expressed in VSMCs play a crucial role in regulating vascular tone. Among these ion channels are CaCCs, Kv and Kir. In VSMCs, Cl^-^ accumulated above the electrochemical equilibrium and when VSMC CaCCs are activated, intracellular Cl^-^ effluxes, leading to the cell membrane depolarization, consequent activation of voltage dependent Ca^2+^ channels and elevation of [Ca^2+^]_i_. Therefore, opening of CaCCs facilitates elevation of the tone of VSMCs [36]. On the other hand, concentration of K^+^ inside VSMCs is much higher than outside. Opening of K^+^ channels leads to hyperpolarization of the cell membrane, consequently depresses [Ca^2+^]_o_ influx and eventually resists the myocyte contraction [37]. Previous studies demonstrated that API relaxed rat aorta and suggested that inhibition of transmembrane [Ca^2+^]_o_ influx through voltage dependent Ca^2+^ channels underlay API-induced vasorelaxation [11,12]. Enhancement of transient receptor potential vanilloid 4 cation channels was also suggested being involved in API-induced vasorelaxation of rat mesenteric arteries [15] and in API-induced protection against hypertension-associated renal damage [38].

To clarify the possible involvement of ion channels other than Ca^2+^ channels in the vascular effects of API, we investigated the impacts of API on CaCCs, Kir and Kv channels in IRA ASMCs using the whole-cell patch clamp technique. As expected in the light of the results of the myograph experiments, at the same concentration range as used in myograph study, API concentration-dependently depressed CaCC currents of IRA ASMCs and shifted the CaCC I-V curves downwards. The depression was reversible because the normal CaCC currents largely recovered after removal of API from the cell bath by washout. As for K^+^ channels, the effects of API were complicated. API enhanced Kv currents while depressed Kir currents of IRA ASMCs. The enhancement on Kv currents is in accordance, while the inhibition on Kir currents is inconsistent with its vasorelaxation in IRAs. This suggests complexity and pleiotropism of API actions. The present study cannot give a satisfactory explanation on this disagreement. In this connection, API actions and the underlying mechanisms on ion channels appeal for further deeper investigation.

Taken together, the present study, for the first time, demonstrated the vasorelaxation of API on IRAs and suggested that depression of CaCCs, enhancement of Kv and increased nitric oxide synthesis may be involved in API-induced IRA relaxation.

## Acknowledgments

This study was supported by the National Natural Science Foundation of China (NSFC 81773738 to MZ, NSFC81603111 to YL) and the Fund for Shanxi “1331 Project” Key Subjects Construction.

## REFERENCES

1. Madunic J, Madunic IV, Gajski G, Popic J, Garaj-Vrhovac V (2018) Apigenin: A dietary flavonoid with diverse anticancer properties. Cancer Lett 413: 11–22.

2. Nabavi SF, Khan H, D’Onofrio G, Samec D, Shirooie S, et al. (2017) Apigenin as neuroprotective agent: Of mice and men. Pharmacol Res.

3. Escande C, Nin V, Price NL, Capellini V, Gomes AP, et al. (2013) Flavonoid apigenin is an inhibitor of the NAD+ ase CD38: implications for cellular NAD+ metabolism, protein acetylation, and treatment of metabolic syndrome. Diabetes 62: 1084–1093.

4. Zhang K, Song W, Li D, Jin X (2017) Apigenin in the regulation of cholesterol metabolism and protection of blood vessels. Exp Ther Med 13: 1719–1724.

5. Ren B, Qin W, Wu F, Wang S, Pan C, et al. (2016) Apigenin and naringenin regulate glucose and lipid metabolism, and ameliorate vascular dysfunction in type 2 diabetic rats. Eur J Pharmacol 773: 13–23.

6. Qin W, Ren B, Wang S, Liang S, He B, et al. (2016) Apigenin and naringenin ameliorate PKCbetaII-associated endothelial dysfunction via regulating ROS/caspase-3 and NO pathway in endothelial cells exposed to high glucose. Vascul Pharmacol 85: 39–49.

7. Jin BH, Qian LB, Chen S, Li J, Wang HP, et al. (2009) Apigenin protects endothelium-dependent relaxation of rat aorta against oxidative stress. Eur J Pharmacol 616: 200–205.

8. Ma X, Li YF, Gao Q, Ye ZG, Lu XJ, et al. (2008) Inhibition of superoxide anion-mediated impairment of endothelium by treatment with luteolin and apigenin in rat mesenteric artery. Life Sci 83: 110–117.

9. Guan H, Gao L, Zhu L, Yan L, Fu M, et al. (2012) Apigenin attenuates neointima formation via suppression of vascular smooth muscle cell phenotypic transformation. J Cell Biochem 113: 1198–1207.

10. Zhang YH, Park YS, Kim TJ, Fang LH, Ahn HY, et al. (2000) Endothelium-dependent vasorelaxant and antiproliferative effects of apigenin. Gen Pharmacol 35: 341–347.

11. Chan EC, Pannangpetch P, Woodman OL (2000) Relaxation to flavones and flavonols in rat isolated thoracic aorta: mechanism of action and structure-activity relationships. J Cardiovasc Pharmacol 35: 326–333.

12. Ko FN, Huang TF, Teng CM (1991) Vasodilatory action mechanisms of apigenin isolated from Apium graveolens in rat thoracic aorta. Biochim Biophys Acta 1115: 69–74.

13. Calderone V, Chericoni S, Martinelli C, Testai L, Nardi A, et al. (2004) Vasorelaxing effects of flavonoids: investigation on the possible involvement of potassium channels. Naunyn Schmiedebergs Arch Pharmacol 370: 290–298.

14. Mastantuono T, Battiloro L, Sabatino L, Chiurazzi M, Di Maro M, et al. (2015) Effects of Citrus Flavonoids Against Microvascular Damage Induced by Hypoperfusion and Reperfusion in Rat Pial Circulation. Microcirculation 22: 378–390.

15. Ma X, He D, Ru X, Chen Y, Cai Y, et al. (2012) Apigenin, a plant-derived flavone, activates transient receptor potential vanilloid 4 cation channel. Br J Pharmacol 166: 349–358.

16. Jackson WF (2005) Potassium channels in the peripheral microcirculation. Microcirculation 12: 113–127.

17. Nelson MT, Quayle JM (1995) Physiological roles and properties of potassium channels in arterial smooth muscle. Am J Physiol 268: C799–822.

18. Betts LC, Kozlowski RZ (2000) Electrophysiological effects of endothelin-1 and their relationship to contraction in rat renal arterial smooth muscle. Br J Pharmacol 130: 787–796.

19. Martens JR, Gelband CH (1996) Alterations in rat interlobar artery membrane potential and K+ channels in genetic and nongenetic hypertension. Circ Res 79: 295–301.

20. Chadha PS, Zunke F, Zhu HL, Davis AJ, Jepps TA, et al. (2012) Reduced KCNQ4-encoded voltage-dependent potassium channel activity underlies impaired beta-adrenoceptor-mediated relaxation of renal arteries in hypertension. Hypertension 59: 877–884.

21. Chilton L, Loutzenhiser R (2001) Functional evidence for an inward rectifier potassium current in rat renal afferent arterioles. Circ Res 88: 152–158.

22. Chilton L, Loutzenhiser K, Morales E, Breaks J, Kargacin GJ, et al. (2008) Inward rectifier K(+) currents and Kir2.1 expression in renal afferent and efferent arterioles. J Am Soc Nephrol 19: 69–76.

23. Matchkov VV, Boedtkjer DM, Aalkjaer C (2015) The role of Ca(2+) activated Cl(-) channels in blood pressure control. Curr Opin Pharmacol 21: 127–137.

24. Liu Y, Niu L, Cui L, Hou X, Li J, et al. (2014) Hesperetin inhibits rat coronary constriction by inhibiting Ca(2+) influx and enhancing voltage-gated K(+) channel currents of the myocytes. Eur J Pharmacol 735: 193–201.

25. Sun H, Xia Y, Paudel O, Yang XR, Sham JS (2012) Chronic hypoxia-induced upregulation of Ca2+-activated Cl-channel in pulmonary arterial myocytes: a mechanism contributing to enhanced vasoreactivity. J Physiol 590: 3507–3521.

26. Cogolludo A, Moreno L, Bosca L, Tamargo J, Perez-Vizcaino F (2003) Thromboxane A2-induced inhibition of voltage-gated K+ channels and pulmonary vasoconstriction: role of protein kinase Czeta. Circ Res 93: 656–663.

27. Wu GB, Zhou EX, Qing DX, Li J (2009) Role of potassium channels in regulation of rat coronary arteriole tone. Eur J Pharmacol 620: 57–62.

28. Park WS, Han J, Kim N, Ko JH, Kim SJ, et al. (2005) Activation of inward rectifier K+ channels by hypoxia in rabbit coronary arterial smooth muscle cells. Am J Physiol Heart Circ Physiol 289: H2461–2467.

29. Tennant BP, Cui Y, Tinker A, Clapp LH (2006) Functional expression of inward rectifier potassium channels in cultured human pulmonary smooth muscle cells: evidence for a major role of Kir2.4 subunits. J Membr Biol 213: 19–29.

30. Park WS, Kim N, Youm JB, Warda M, Ko JH, et al. (2006) Angiotensin II inhibits inward rectifier K+ channels in rabbit coronary arterial smooth muscle cells through protein kinase Calpha. Biochem Biophys Res Commun 341: 728–735.

31. Smith PD, Brett SE, Luykenaar KD, Sandow SL, Marrelli SP, et al. (2008) KIR channels function as electrical amplifiers in rat vascular smooth muscle. J Physiol 586: 1147–1160.

32. Chilton L, Smirnov SV, Loutzenhiser K, Wang X, Loutzenhiser R (2011) Segment-specific differences in the inward rectifier K(+) current along the renal interlobular artery. Cardiovasc Res 92: 169–177.

33. Hayoz S, Pettis J, Bradley V, Segal SS, Jackson WF (2017) Increased amplitude of inward rectifier K(+) currents with advanced age in smooth muscle cells of murine superior epigastric arteries. Am J Physiol Heart Circ Physiol 312: H1203–H1214.

34. Elhennawy MG, Lin HS (2017) Quantification of apigenin trimethyl ether in rat plasma by liquid chromatography-tandem mass spectrometry: Application to a pre-clinical pharmacokinetic study. J Pharm Biomed Anal 142: 35–41.

35. Zhang J, Huang Y, Liu D, Gao Y, Qian S (2013) Preparation of apigenin nanocrystals using supercritical antisolvent process for dissolution and bioavailability enhancement. Eur J Pharm Sci 48: 740–747.

36. Hubner CA, Schroeder BC, Ehmke H (2015) Regulation of vascular tone and arterial blood pressure: role of chloride transport in vascular smooth muscle. Pflugers Arch 467: 605–614.

37. Salomonsson M, Brasen JC, Sorensen CM (2017) Role of renal vascular potassium channels in physiology and pathophysiology. Acta Physiol (Oxf).

38. Wei X, Gao P, Pu Y, Li Q, Yang T, et al. (2017) Activation of TRPV4 by dietary apigenin antagonizes renal fibrosis in deoxycorticosterone acetate (DOCA)-salt-induced hypertension. Clin Sci (Lond) 131: 567–581.

